# Purkinje cell dysfunction causes disrupted sleep in ataxic mice

**DOI:** 10.1101/2023.07.03.547586

**Authors:** Luis E. Salazar Leon, Amanda M. Brown, Heet Kaku, Roy V. Sillitoe

**Affiliations:** Department of Neuroscience, Baylor College of Medicine, Houston, Texas, USA; Department of Pathology & Immunology, Baylor College of Medicine, Houston, Texas, USA; Jan and Dan Duncan Neurological Research Institute at Texas Children’s Hospital, Houston, Texas, 77030, USA; Department of Pediatrics, Baylor College of Medicine, Houston, Texas, USA; Development, Disease Models & Therapeutics Graduate Program, Baylor College of Medicine, Houston, Texas, USA

**Keywords:** Purkinje cells, cerebellar nuclei, ataxia, sleep, circadian rhythms

## Abstract

Purkinje cell dysfunction causes movement disorders such as ataxia, however, recent evidence suggests that Purkinje cell dysfunction may also alter sleep regulation. Here, we used an ataxia mouse model generated by silencing Purkinje cell neurotransmission (*L7^Cre^;Vgat^fx/fx^*) to better understand how cerebellar dysfunction impacts sleep physiology. We focused our analysis on sleep architecture and electrocorticography (ECoG) patterns based on their relevance to extracting physiological measurements during sleep. We found that circadian activity is unaltered in the mutant mice, although their sleep parameters and ECoG patterns are modified. The *L7^Cre^;Vgat^fx/fx^* mutant mice have decreased wakefulness and rapid eye movement (REM) sleep, while non-rapid eye movement (NREM) sleep is increased. The mutant mice have an extended latency to REM sleep, which is also observed in human ataxia patients. Spectral analysis of ECoG signals revealed alterations in the power distribution across different frequency bands defining sleep. Therefore, Purkinje cell dysfunction may influence wakefulness and equilibrium of distinct sleep stages in ataxia. Our findings posit a connection between cerebellar dysfunction and disrupted sleep and underscore the importance of examining cerebellar circuit function in sleep disorders.

**Summary Statement:** Utilizing a precise genetic mouse model of ataxia, we provide insights into the cerebellum’s role in sleep regulation, highlighting its potential as a therapeutic target for motor disorders-related sleep disruptions.

## Introduction

The cerebellum is critical for different motor functions including coordination, posture, balance, and learning. However, emerging evidence overwhelmingly suggests that the cerebellum also plays a crucial role in non-motor functions, ranging from cognitive and emotional processing(Popa et al., 2014), associative learning(Larry et al., n.d.), to reward expectation(Carta et al., 2019). Recent work suggests that the cerebellum also plays a role in the regulation of sleep and sleep-associated processes(Salazar Leon and Sillitoe, 2022; Song and Zhu, 2021; Strick et al., 2009). Indeed, the association between cerebellar dysfunction and sleep disturbances has been corroborated in both human patients and recently in mouse models with dystonia(Leon and Sillitoe, 2023). In humans, sleep anomalies primarily manifest as disruptions in sleep timing, resulting in daytime drowsiness, increased sleep latency denoting difficulties initiating sleep, and parasomnia reflecting problems with sleep maintenance(Antelmi et al., 2017; Smit et al., 2017). Importantly, these deficits disproportionately impact rapid eye movement (REM) sleep. Except for daytime drowsiness, analogous impairments are observable in mouse models of dystonia, particularly mirroring human deficits in REM sleep timing and latency(Leon and Sillitoe, 2023). A similar association has been observed in human patients with spinocerebellar ataxia, a neurodegenerative motor disorder characterized by uncoordinated movements; patients with ataxia present with substantial sleep disruptions, similar to patients with dystonia(Huebra et al., 2019; Pedroso et al., 2011; Shindo et al., 2019). In particular, and again similarly to human patients with dystonia, sleep disruptions in ataxia tend to involve specific impairments in REM sleep length and quality. For ataxia patients, the severity of ataxia is also a predictor of sleep impairments(Sonni et al., 2014).

While sleep impairments in motor diseases are typically considered secondary symptoms, dysfunction in normal sleep behavior substantially impacts patient health. Studies in patients with cerebellar ataxia reveal that cognitive function and depression are equally and positively correlated with sleep quality(Sonni et al., 2014). Disrupted sleep also has known associations with impairments in motor learning and motor function, including eyeblink conditioning (in mice) and finger-tapping tasks (in humans) as well as motor symptoms in patients with motor diseases(De Zeeuw and Canto, 2020; Walker et al., 2003). It is postulated that this association results from the modulation of synaptic activity during sleep(Tononi and Cirelli, 2014). Thus, it stands to reason that sleep may have a key role in mediating motor function in the context of motor diseases like cerebellar ataxia. However, while the cerebellum has many known links to abnormal motor function, the cerebellar circuit components involved in sleep regulation remain unclear. This problem is magnified in the context of ataxia, where cerebellar function is directly affected.

Of particular interest is the role of Purkinje cells, the sole output neurons of the cerebellar cortex. Purkinje cells have been established as a central driver of the motor phenotypes of ataxia, both in animal models(White et al., 2014) and human patients(Xia et al., 2013). Indeed, in many models of ataxia the disease onset and progression are driven by Purkinje cell signaling deficits(Hoxha et al., 2018; White et al., 2014). Interestingly, Purkinje cell signals are also known to demonstrate sleep-dependent activity, increasing their firing during NREM sleep and prior to the transition between sleep and wake(Mano, 1970; Zhang et al., 2020). Despite the growing body of literature linking cerebellar dysfunction to sleep disruption, the specific consequences of Purkinje cell dysfunction on sleep regulation remain unclear. Prior research conducted in our laboratory has substantiated that disruptions in olivocerebellar signaling in mice result in both motor (dystonic) and nonmotor (sleep) dysfunction(Leon and Sillitoe, 2023). While this observation served as robust evidence suggesting that the cerebellum may independently precipitate motor and sleep dysfunctions, the precise extent to which direct manipulation of cerebellar circuits could affect sleep remained ambiguous, as the genetic modifications in the previous work primarily focused on silencing inputs into the cerebellum rather than neurotransmission of the cerebellar circuit itself. Considering the critical role of Purkinje cells, which function as the primary integrators and processors of cerebellar inputs, we aimed to explore whether these cells, and by extension the cerebellum, could be a primary driving force behind sleep impairments. Therefore, based on our recent work, we propose here that cerebellar dysfunction might instigate sleep impairments across different states of disease.

The central problem that guided this inquiry was whether direct cerebellar manipulation, specifically through modulating Purkinje cell activity, could lead to measurable changes in sleep patterns. Therefore, the focus of our current study extended beyond the input pathways to the cerebellum, to the processing and output stages within the cerebellum itself, with an emphasis on the potential role of Purkinje cells in mediating both movement and sleep disturbances. The primary motivation of this work was to clarify the role of cerebellar circuitry in sleep regulation, and by doing so, potentially highlight the underlying mechanisms that may contribute to the sleep impairments in cerebellar disorders.

To investigate this relationship between Purkinje cell function and sleep, we used a constitutively active Cre-loxP system to block inhibitory synaptic transmission of Purkinje cells. As Purkinje cells are inhibitory neurons and represent the sole output of the cerebellar cortex, this genetic manipulation effectively eliminates communication between the cerebellar cortex and the downstream cerebellar nuclei by blocking fast neurotransmission. *Vgat* was deleted using a *Pcp2* (*L7*) gene regulatory element to spatially drive and restrict Cre expression to Purkinje cells. The resulting mice had the genotype *L7^Cre^;Vgat^fx/fx^*and presented with severe ataxic motor symptoms, including widened gait, incoordination, and a lack of balance. This ataxia mouse model (*L7^Cre^;Vgat^fx/fx^*) was previously generated by our lab and its ataxic motor deficits have been explored in detail(White et al., 2014). Due to the severe and reproducible ataxic phenotype of *L7^Cre^;Vgat^fx/fx^*mice, we were presented with an opportunity to explore sleep processes in the context of cerebellar ataxia. Furthermore, due to the highly restricted nature of our genetic manipulation that targets Purkinje cells, we could specifically test the interaction between cerebellar dysfunction and sleep.

In this work, we report that *L7^Cre^;Vgat^fx/fx^*mice display significant sleep impairments, some of which mirror those observed in human patients with cerebellar ataxia. Although circadian activity remains unaltered in the mutants, they do exhibit modified sleep parameters and distinctly different ECoG patterns compared to control mice. *L7^Cre^;Vgat^fx/fx^* mice demonstrate decreased wakefulness and REM sleep, while NREM sleep is increased. The mutant mice also show an extended latency to REM sleep, a result congruent with existing research on sleep quality in human ataxia patients(Sonni et al., 2014). Additionally, spectral analysis of ECoG signals reveals alterations in the power distribution across different frequency bands that define sleep, across both frontal and parietal cortices, further indicative of disrupted mechanisms of sleep homeostasis. Our work suggests that Purkinje cell dysfunction may influence not only wakefulness but also the equilibrium of distinct sleep stages, potentially contributing to sleep-related complications in cerebellar disorders like ataxia. Our findings underscore the importance of exploring cerebellar involvement in sleep regulation, and how Purkinje cells may serve as key regulators of sleep-wake rhythms in the mammalian brain.

## Results

### Purkinje cell silencing in *L7^Cre^;Vgat^fx/fx^* mice occurs throughout the cerebellum

In previous work we demonstrated that genetically silencing Purkinje cell gamma-Aminobutyric acid (GABA) neurotransmission can induce severe ataxic motor phenotypes in mice(White et al., 2014). The *L7^Cre^;Vgat^fx/fx^*mouse model utilizes the Cre-loxP system to drive the deletion of *Vgat* in neurons in which the *L7* promoter is active (Figure 1A). In a control (*Vgat^fx/fx^*) mouse, normal Purkinje cell function is maintained; when an action potential reaches an axon terminal, the vesicular GABA transporter (VGAT) supports the loading of GABA into pre-synaptic vesicles. This process enables the action potential to propagate as GABA traverses the synaptic cleft, effectively facilitating fast neurotransmission. Anatomically, inhibitory efferents project from the Purkinje cell layer within the cerebellar cortex, subsequently synapsing with the cerebellar nuclei neurons, the primary output cells of the cerebellum (Figure 1B). This means that, in *L7^Cre^;Vgat^fx/fx^*mice, the loss of Purkinje cell-specific *Vgat* results in the silencing of Purkinje cell-to-cerebellar nuclei neuron neurotransmission, as GABA cannot be loaded onto vesicles in the absence of VGAT (Figure 1A, 1C). We validated that *L7*-mediated *Cre* expression in *L7^Cre^;Vgat^fx/fx^* mice was present throughout the cerebellar cortex (Figure 1D), which resulted in a complete loss of *Vgat* expression in Purkinje cells (Figure 1F). In *L7^Cre^;Vgat^+/+^* control mice, *Vgat* expression is typically highly co-localized with *Cre* expression in the Purkinje cell layer of the cerebellar cortex. In these control mice, no deletion of *Vgat* occurs (Figure 1E). Other cerebellar neuron types are present in their normal anatomical locations despite this manipulation(White et al., 2014). These findings confirm that the previously characterized ataxic phenotypes (impaired balance, lack of motor coordination, wide-based gait) are a result of silenced cerebellar Purkinje cells(Brown et al., 2020; White et al., 2014) and further support the use of this animal model to specifically probe the role of cerebellar Purkinje cells in regulating sleep.

**Figure 1:**
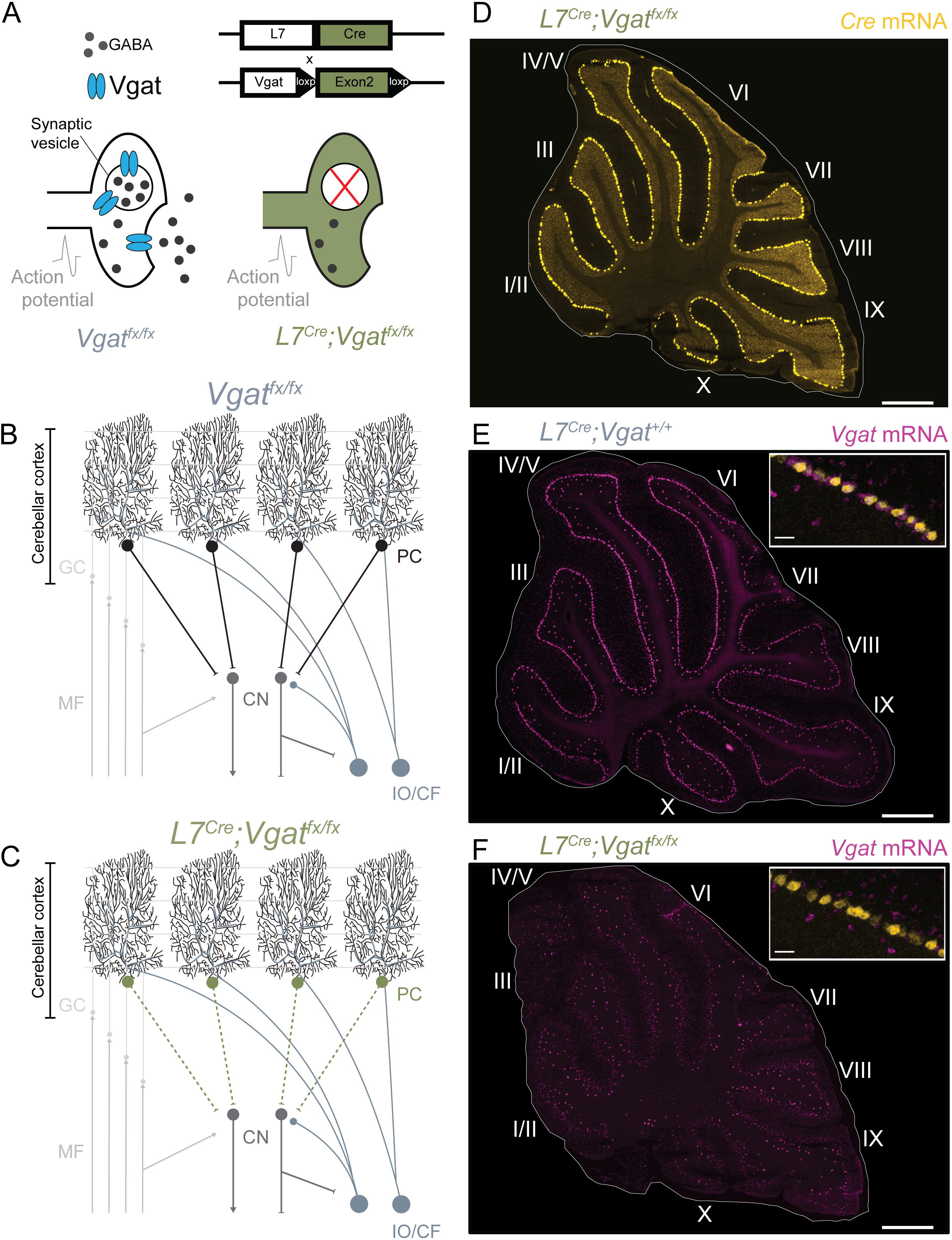
*L7^Cre^;Vgat^fx/fx^* mice display *Vgat* deletion throughout the cerebellar cortex. **(A)** Using the *L7^Cre^* genetic driver line, exon 2 of *Vgat* was selectively removed from Cre-expressing cells. This resulted in deletion of VGAT expression with spatial sensitivity, and subsequent silencing in the affected cells. **(B)** Schematic demonstrating a simplified cerebellar circuit in *Vgat^fx/fx^* control mice, in which all components of cerebellar signaling are intact and functional. Abbreviations: MF, mossy fiber; GC, granule cell; PC, Purkinje cell; CN, cerebellar nuclei; IO/CF, inferior olive/climbing fiber. **(C)** Schematic demonstrating the result of *Vgat* deletion in *L7^Cre^;Vgat^fx/fx^* mutant mice. Mice display widespread silencing of Purkinje cell synaptic output. Abbreviations: same as **(B)**. **(D)** Image processed using *in situ* hybridization showing widespread *L7*-mediated *Cre* expression throughout the cerebellar cortex on adult mouse sagittal cerebellar sections in an *L7^Cre^;Vgat^fx/fx^* mutant mouse. **(E)** Same as **(D)** but for *Vgat* expression in a control mouse expressing *L7Cre* but with no floxed copies of *Vgat*. The inset shows co-expression of *Cre* and *Vgat* in the cerebellar cortex. **(F)** Same as **(E)** but for an *L7^Cre^;Vgat^fx/fx^* mutant mouse. Inset shows co-expression of *Cre* and *Vgat*, and lack of *Vgat* expression as expected.

### Circadian activity is unchanged in *L7^Cre^;Vgat^fx/fx^* mutant mice

Sleep regulation is thought to be governed by two processes: the homeostatic process (Process S), which reflects the build-up of sleep pressure during wakefulness, and the circadian process (Process C), which regulates the timing of sleep and wakefulness based on the 24-hour biological clock(Borbély, 2022) (Figure 2A). Many methods exist for assessing the strength and stability of Process C. Wheel-running activity is commonly used in rodents as a non-invasive proxy for measuring daily activity patterns(Eckel-Mahan and Sassone-Corsi, 2015), and previous work has used wheel-running to assess circadian activity in other mouse models of mild ataxia(Mendoza et al., 2010) and dystonia(Leon and Sillitoe, 2023). As human ataxia patients typically present with circadian deficits like fatigue and excessive daytime sleepiness alongside changes in sleep quality(Sonni et al., 2014), we sought to determine whether circadian activity was similarly affected in *L7^Cre^;Vgat^fx/fx^* mutant mice. Mice were singly housed with *ad libitum* access to food, water, and a running wheel within their home cage (Figure 2B). Revolutions of the running wheels were automatically monitored for the entire duration of the recording period (14 days baseline (LD; light-dark), 21 days constant condition (DD; dark-dark) Figure 2C). Data were collected and plotted as actograms, in which each row represents a day and black tick marks represent revolutions of the running wheel, indicative of locomotor activity, and therefore are interpreted as a period of wake. The data was double-plotted such that 48 hours of activity is represented on a single line, to better visualize the patterns of activity(Eckel-Mahan and Sassone-Corsi, 2015). We observed normal nocturnal behavior for *Vgat^fx/fx^* control mice for the 14-day baseline LD period, followed by typically free-running behavior in the DD period (Figure 2D). Given their severe ataxic phenotype, we predicted that wheel-running behavior in *L7^Cre^;Vgat^fx/fx^* mice would be limited, and indeed found that mutant mice required significant time to begin running consistently. Interestingly, we did observe some activity that was qualitatively normal compared to controls, particularly during the DD period (Figure 2E). As expected, average activity counts for *L7^Cre^;Vgat^fx/fx^* mutant mice were significantly lower during both LD and DD periods (Figure 2F, 2G). Average period length (the time it takes for a circadian rhythm to complete a full cycle) during the DD condition was not significantly different between mutants and controls, suggesting that endogenous circadian activity remained intact (Figure 2H). We also measured onset phase shift, which is the change in the timing of the onset of activity (the beginning of the active phase) in response to a manipulation. Here, the manipulation is the change from LD to DD conditions, and onset phase shift measures the gradual shift in the timing of activity onset from day 14 (DD start) until day 35 (experiment end). We found that onset phase shift was similar between *L7^Cre^;Vgat^fx/fx^* mutant mice and controls (Figure 2I). These results suggest that circadian activity remains unchanged in *L7^Cre^;Vgat^fx/fx^* mutant mice, despite their cerebellar and motor dysfunction.

**Figure 2:**
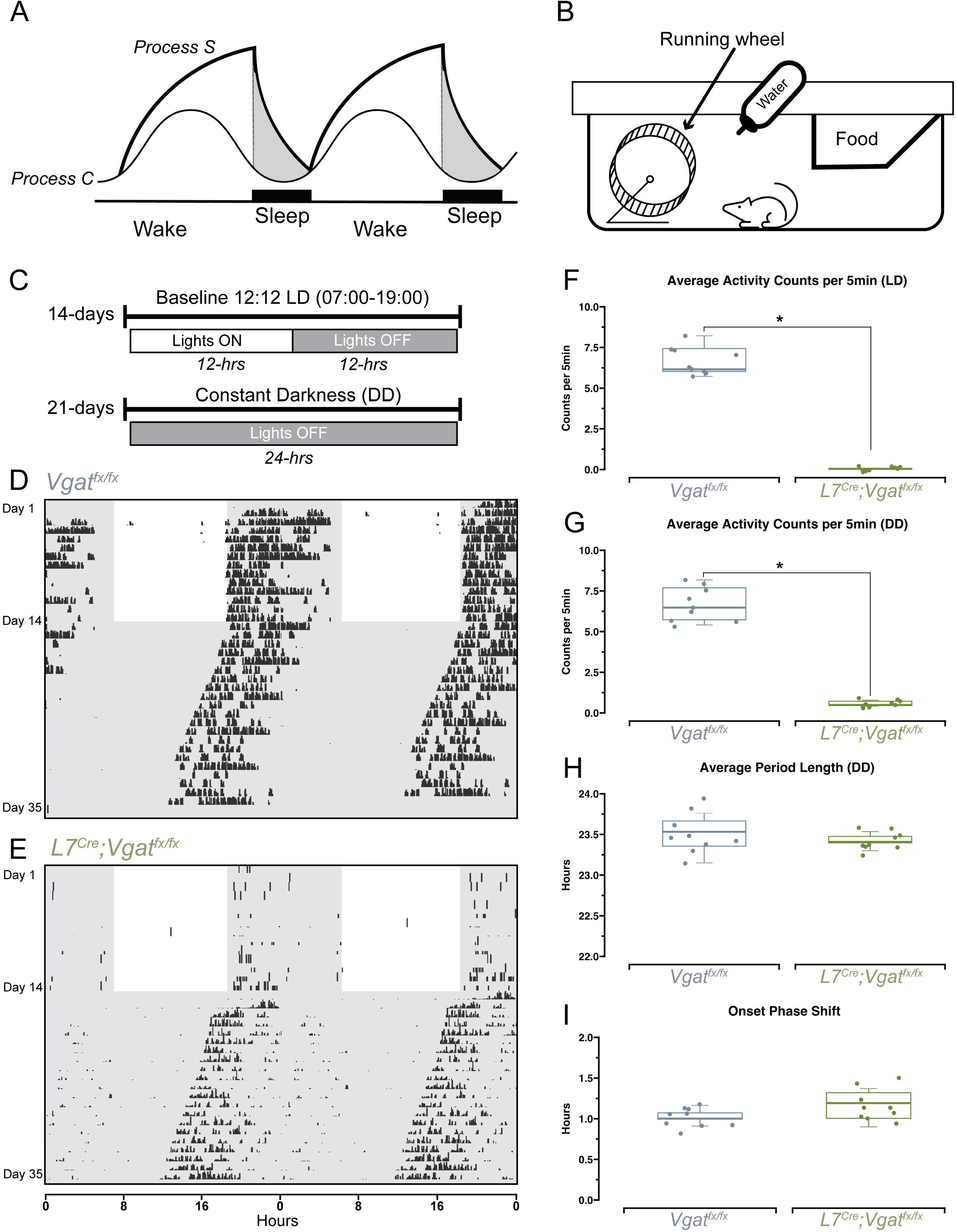
Circadian wheel-running behavior is normal in *L7^Cre^;Vgat^fx/fx^* mutant mice. **(A)** A schematic representation of the two-process model of sleep regulation. Process S denotes the homeostatic drive to sleep, Process C denotes the circadian drive to sleep. **(B)** Schematic illustration of the home cage setup for wheel-running. **(C)** Timeline of the wheel-running experiment. Light-dark (LD), dark-dark (DD). **(D)** Representative double-plotted actogram for a control mouse. Rows represent days, black ticks represent revolutions of the wheel, indicative of locomotor activity. Unshaded regions represent lights on, black shaded regions represent lights off. **(E)** Same as **(D)** but for an *L7^Cre^;Vgat^fx/fx^* mutant mouse. **(F)** Quantification of average activity counts per 5 minutes, only during the light-dark (LD) phase. **(G)** Same as **(F)** but for the dark-dark (DD) phase. **(H)** Quantification of average period length, only during the DD phase. **(I)** Quantification of onset phase shift, from day 14 to day 35 of recording. Points on **F-I** represent individual mice (n=9 per group). All source data and specific p-values are available in Figure 2 – source data 1.

### Ataxic *L7^Cre^;Vgat^fx/fx^* mice display significantly disrupted stages of sleep

The relationship between sleep and cerebellar ataxia is particularly relevant. Patients with ataxia not only display disrupted sleep(Patterson et al., 2018; Shindo et al., 2019; Sonni et al., 2014), but also ataxic symptom severity is directly associated with sleep dysfunction. The combination subsequently has a direct impact on additional factors that affect quality of life, including fatigue and depression(Patterson et al., 2018; Sonni et al., 2014). Therefore, a major goal was to determine the sleep architecture of *L7^Cre^;Vgat^fx/fx^*mice. We therefore implanted *L7^Cre^;Vgat^fx/fx^* mutants and *Vgat^fx/fx^* controls with platinum-iridium ECoG and EMG electrodes and recorded signals continuously during the light-phase, when mice naturally sleep (Figure 3 A-C). ECoG/EMG waveforms show that *L7^Cre^;Vgat^fx/fx^* mice display the typical spectral activity, which defines the arousal states of wake, REM sleep, and NREM sleep (Figure 3D). We then assessed the total time spent in each arousal state, for the duration of the recording period. We note that while typical sleep cycles in mice are shorter than in humans, they still follow the same general pattern of wake, followed by NREM sleep, followed by REM sleep (Figure 3E). Hypnograms from 1 hour of the recording period suggest that *L7^Cre^;Vgat^fx/fx^* mutants display disrupted sleep patterns relative to controls. Periods of wake were less frequent while the length of individual bouts of NREM sleep were extended (Figure 3F). Upon analyzing the proportions of time spent in each state for the entire recording period, we found that *L7^Cre^;Vgat^fx/fx^* mice spent significantly less time awake, less time in REM, and greater time in NREM sleep (Figure 3 G-I). These results suggest that the Purkinje cell-specific manipulation that alters activity in *L7^Cre^;Vgat^fx/fx^*mice is sufficient to drive impairments in sleep patterns, further supporting a role for the cerebellum in regulating sleep.

**Figure 3:**
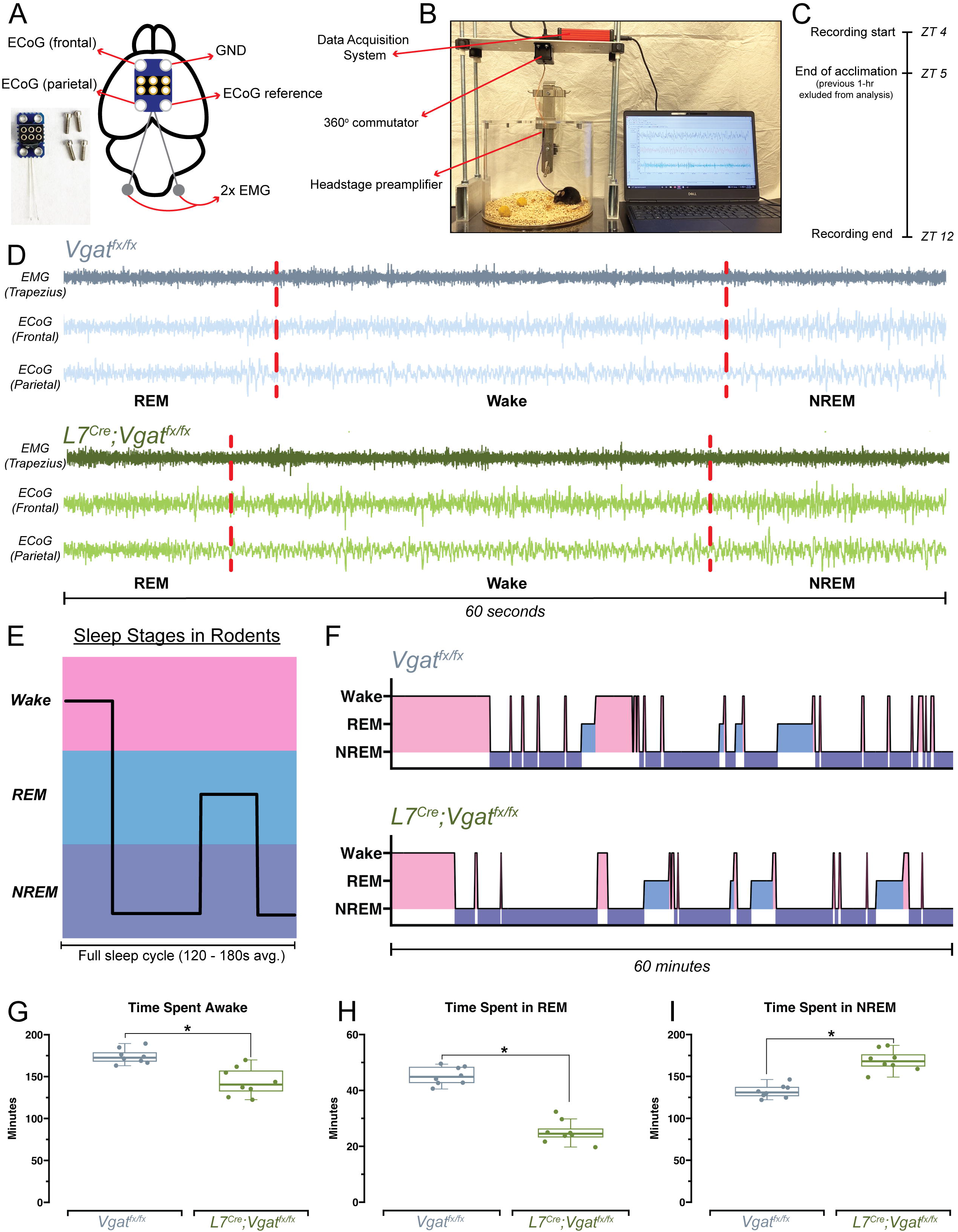
Sleep patterns are disrupted in *L7^Cre^;Vgat^fx/fx^* mutant mice. **(A)** Schematic illustration of a mouse brain, with ECoG/EMG headmount electrode placement. An image of the ECoG/EMG electrode and recording screws is also shown in the bottom left. **(B)** An image of the ECoG/EMG sleep recording setup. **(C)** A schematic of the experimental timeline for recording sleep from each mouse. **(D)** Raw waveforms of EMG (top trace) and ECoG (bottom traces) recorded from a control and *L7^Cre^;Vgat^fx/fx^*mutant mouse. Example sample trace is 60 seconds in length. Sleep stage is noted at the bottom of each trace and differentiated by dashed red lines. **(E)** Schematic demonstrating sleep stages and their relative depth and temporal organization in mice. **(F)** Hypnograms for one representative *Vgat^fx/fx^* mouse (control, top) and *L7^Cre^;Vgat^fx/fx^*mouse (mutant, bottom). Both hypnograms refer to the same 1-hour period, from 1PM to 2PM CST. Periods of wake, REM, and NREM are highlighted and correspond to the example schematic in **(E)**. **(G)** Quantification of total time spent awake. **(H)** Quantification of total time spent in REM. **(I)** Quantification of total time spent in NREM. Points on **G-I** represent individual mice (n=8 per group). All source data and specific p-values are available in Figure 3 – source data 1.

### Lack of Purkinje cell neurotransmission causes sleep pattern disruptions that can be defined by enhanced NREM at the expense of wake and REM phases

We observed that the Purkinje cell-initiated cerebellar ataxia in *L7^Cre^;Vgat^fx/fx^* mice was sufficient to disrupt sleep stages, with a particular impact on REM sleep. This is intriguingly similar to human patients with ataxia, whose sleep disruptions disproportionally affect the timing and quality of REM sleep(Patterson et al., 2018; Pedroso et al., 2011; Sonni et al., 2014). Still, it is unclear how these patterns of disruption arise, and whether changes in frequency or length of sleep stages (or both) are responsible for driving the observed sleep impairments (Figure 4 A-C). Therefore, to further understand the specific disruption of each stage, we assessed the total number of arousal-state bouts and the average length of each bout for wake, REM, and NREM sleep. All calculations were performed after the defined onset of sleep, which was determined according to previous work(Hunsley and Palmiter, 2004; Leon and Sillitoe, 2023). Consistent with our initial findings of sleep disruption, we found that the total number of wake bouts was significantly lower in *L7^Cre^;Vgat^fx/fx^* mice (Figure 4D). Similarly, the total number of REM bouts was decreased while the total number of NREM bouts was increased (Figure 4E, 4F). Interestingly, we found that despite their overall decreased occurrence, the average length of wake bouts was longer for *L7^Cre^;Vgat^fx/fx^*mutant mice, suggesting the presence of a barrier to falling asleep or a reduced sleep pressure (Figure 4G). We also found that the average length of both REM and NREM bouts was longer in *L7^Cre^;Vgat^fx/fx^* mice (Figure 4H, 4I). Previous work in human patients suggests a deficit not only in the quantity of REM sleep but also the time to initially achieve REM sleep(Sonni et al., 2014) (REM latency). Given our results of decreased REM time, we hypothesized that a similar deficit for REM latency may exist in the *L7^Cre^;Vgat^fx/fx^*mice. To this end, we calculated the latency to reach both REM and NREM sleep, to assess whether sleep disruption in *L7^Cre^;Vgat^fx/fx^* mice is primarily related to difficulties in falling asleep or staying asleep (or both) (Figure 4J). Both the *L7^Cre^;Vgat^fx/fx^*mice and the *Vgat^fx/fx^* controls displayed similar NREM latency (Figure 4L). However, the *L7^Cre^;Vgat^fx/fx^* mice had significantly elevated latency to REM sleep, by over 1 hour (Figure 4K). Together, these experiments define the specific sleep deficits in *L7^Cre^;Vgat^fx/fx^* mice as an overall increase in NREM time due to an increase in number and duration of NREM bouts at the expense of time spent in both wake and REM. While there is an increase in the duration of wake and REM bouts, this increase in duration cannot overcome the overall reduction in number of wake and REM bouts, resulting in an overall reduction of time spent awake and in REM. The deficits in REM extend to the latency to achieve REM, which is prolonged in *L7^Cre^;Vgat^fx/fx^* mice. These experiments further highlight the degree to which the quality, quantity, and overall timing of wake and sleep phases are dependent on Purkinje cell GABA neurotransmission and support the occurrence of REM sleep deficits in ataxia.

**Figure 4:**
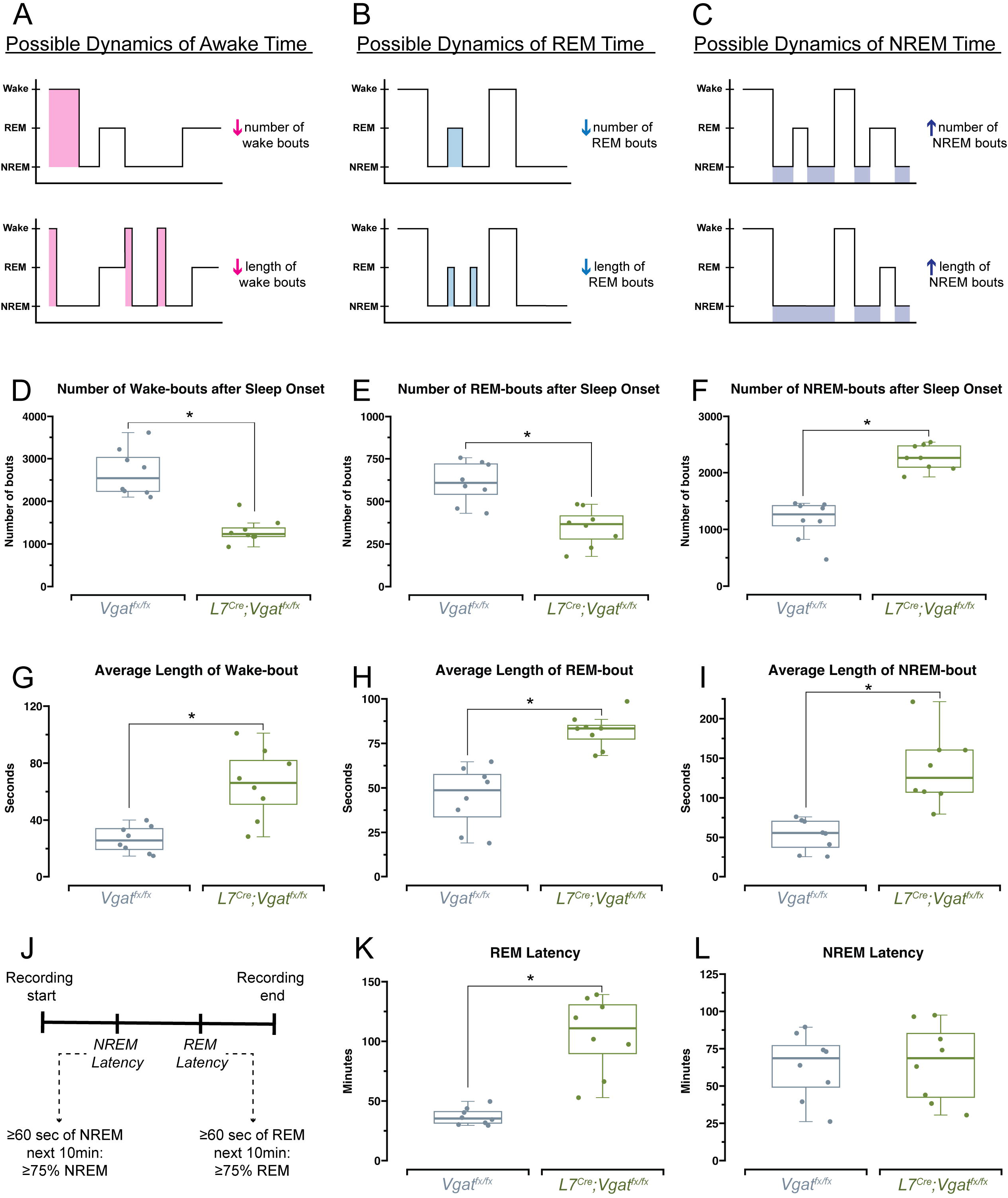
*L7^Cre^;Vgat^fx/fx^* mice have impairments in the quantity and length of sleep bouts. **(A)** Schematic representation of hypnograms showing two hypothesized explanations for changes in awake time, affecting number of bouts or length of bouts. **(B)** Same as **(A)** but for REM time. **(C)** Same as **(A)** but for NREM time. **(D)** Quantification of the number of awake bouts after sleep onset. **(E)** Quantification of the number of REM bouts after sleep onset. **(F)** Quantification of the number of NREM bouts after sleep onset. **(G)** Quantification of the average length of awake bouts. **(H)** Quantification of the average length of REM bouts. **(I)** Quantification of the average length of NREM bouts. **(J)** Schematic showing how REM and NREM latency were calculated, as in Hunsley & Palmiter, 2004 (Hunsley and Palmiter, 2004). **(K)** Quantification of latency to REM sleep. **(L)** Quantification of latency to NREM sleep. Points on **D-I, K, L,** represent individual mice (n=8 per group). All source data and specific p-values are available in Figure 4 – source data 1.

### Sleep state impairments in *L7^Cre^;Vgat^fx/fx^* mice correspond to alterations in delta, theta, alpha, beta, and gamma frequency bands across the frontal and parietal cortices

Wake, REM, and NREM arousal states are defined by specific spectral frequency oscillations, which occur at frequency bands ranging from 0.5 to >100 Hz (Figure 5A, 5B). Transitions between sleep stages in mice can be described in part by changes in delta (0.5-4 Hz), theta (5-8 Hz), and alpha (8-13 Hz) frequency bands, which primarily correspond to NREM, REM, and awake arousal states respectively(BJORNESS et al., 2018; Long et al., 2021). However, changes in higher frequency bands including beta (13-30 Hz) and gamma (35-44 Hz) frequency bands can also indicate not only disruptions in other neuronal processes like associative memory consolidation or sensory processing(Posada-Quintero et al., 2019), but may also play a role in sleep-specific processes such as the maintenance of sleep homeostasis(Grønli et al., 2016). In this way, understanding changes in spectral frequency oscillations can help to better frame changes in sleep-wake dynamics, particularly as different frequency bands can be used to report overall changes in brain connectivity(Torres-Herraez et al., 2022). We therefore determined whether *L7^Cre^;Vgat^fx/fx^*mice displayed quantifiable differences in spectral frequency oscillations in delta, theta, alpha, beta, and gamma frequency bands, across both recording electrodes (Figure 5B). The same cortical (ECoG) electrodes used to record sleep states were used to detect changes in oscillation spectral power frequency throughout the recording period. The two independent ECoG electrodes were placed above the frontal and parietal cortices, and average spectral power from each region, for each frequency band of interest, was assessed. The *L7^Cre^;Vgat^fx/fx^*mice displayed significantly elevated delta power as measured over parietal cortex, but not frontal cortex (p=0.05) (Figure 5C, 5D). Theta (Figure 5E, 5F), alpha (Figure 5G, 5H) and beta (Figure 5I, 5J) power were significantly decreased in the *L7^Cre^;Vgat^fx/fx^*mice, but only in the parietal cortex. In contrast, gamma power was decreased in *L7^Cre^;Vgat^fx/fx^* mice only in frontal but not parietal cortex (Figure 5K, 5L). These data demonstrate that *L7^Cre^;Vgat^fx/fx^* mutants and *Vgat^fx/fx^* controls have measurable differences in spectral frequency bands relevant for sleep, and that these differences are not homogenous throughout the cortex. Additionally, the dynamics of these changes in spectral activity concur with the directionality of overall changes in wake, REM, and NREM time, further reinforcing that they are a direct result of our cerebellar manipulation.

**Figure 5:**
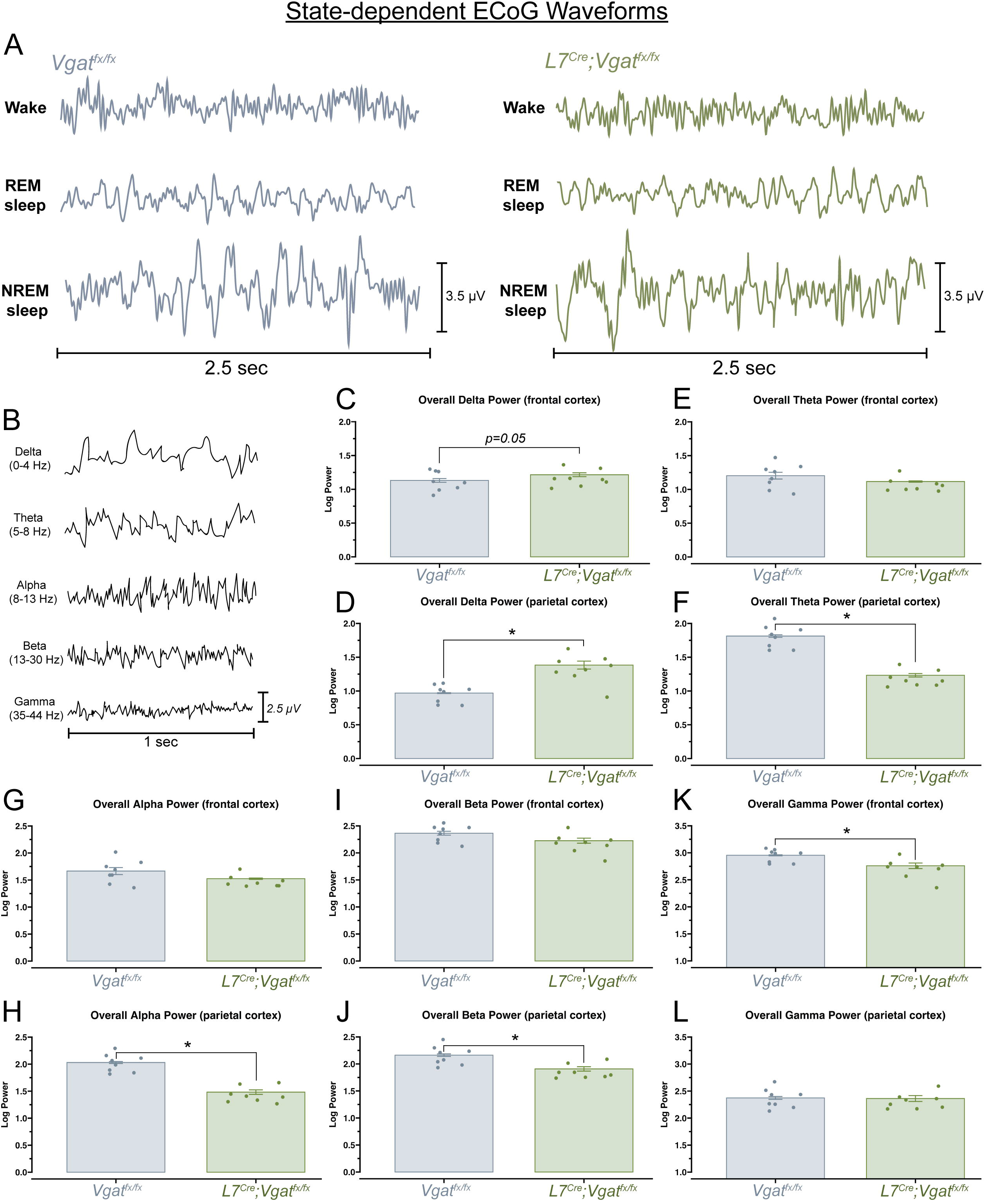
Changes in delta, theta, alpha, beta, and gamma frequency bands accompany sleep impairments in *L7^Cre^;Vgat^fx/fx^* mice. **(A)** 2.5 second examples of raw ECoG waveforms of wake, REM, and NREM sleep from a *Vgat^fx/fx^* (control) and *L7^Cre^;Vgat^fx/fx^*(mutant) mouse. **(B)** 1 second examples of raw ECoG waveforms from a *Vgat^fx/fx^* control mouse, for each frequency band of interest. **(C)** Quantification of overall delta power (0-4 Hz) over frontal cortex. Average power across entire recording period. **(D)** Same as **(C)** but from parietal cortex. **(E)** Quantification of overall theta power (5-8 Hz) over frontal cortex. Average power across entire recording period. **(F)** Same as **(E)** but from parietal cortex. **(G)** Quantification of overall alpha power (8-13 Hz) over frontal cortex. Average power across entire recording period. **(H)** Same as **(G)** but from parietal cortex. **(I)** Quantification of overall beta power (13-30 Hz) over frontal cortex. Average power across entire recording period. **(J** Same as **(I)** but from parietal cortex. **(K)** Quantification of overall gamma power (35-44 Hz) over frontal cortex. Average power across entire recording period. **(L)** Same as **(K)** but from parietal cortex. Points on **C-L** represent individual mice (n=8 per group). All source data and specific p-values are available in Figure 5 – source data 1.

## Discussion

In this study, we show that a genetic manipulation, which precisely targets one cerebellar cell type and results in motor deficits, additionally causes perturbations in the dynamics of arousal states. We utilized a mouse model (*L7^Cre^;Vgat^fx/fx^*) that manifests with Purkinje cell dysfunction and ataxia, thereby offering a platform to scrutinize the cerebellum’s contribution to sleep disturbances with relevance to motor disease. We demonstrate that blocking Purkinje cell GABA neurotransmission is uniform across the cerebellar cortex (Figure 1D-F). While this manipulation resulted in severe ataxia that reduced overall movement (Figure 2F-G), it did not affect the underlying circadian activity (Figure 2D-E, 2H-I). However, the proportion of arousal states and patterning of the phases of sleep were severely disrupted in mice lacking Purkinje cell neurotransmission (Figure 3F-I, 4D-L). This was further evidenced by alterations in ECoG frequency band powers in the frontal and parietal cortices (Figure 5). Therefore, our work suggests that cerebellar Purkinje cells are necessary for normal sleep patterning and contribute to its disruption in the context of ataxia.

Our previous work on manipulating cerebellar inputs that cause dystonia and sleep deficits(Leon and Sillitoe, 2023), combined with the results of the current work, provide evidence for the role of the cerebellum in mediating aspects of sleep dysfunction, particularly in the context of various motor disorders (Figure 6A). The contrast between our previous work, which silenced cerebellar inputs, and the current study, which inhibits Purkinje cell output, offers intriguing insights into the complex cerebellar circuitry governing sleep. Notably, despite the different target within the cerebellar circuit, neither manipulation influenced circadian rhythms of activity; however, both led to substantive sleep impairments, specifically impacting the quality, timing, and frequency of wake, NREM, and REM sleep. Both models resulted in decreased total time in REM with increased total time in NREM. The primary difference remains that, in our prior animal models of dystonia, where cerebellar inputs at the olivocerebellar synapses were silenced, there was a global increase in wakefulness. Conversely, in the present study, mice with silenced Purkinje cells exhibited global reductions in awake time. Therefore, considering these findings, our combined results highlight the pivotal role of proper Purkinje cell activity in regulating sleep-wake cycles. This reinforces the notion that, despite the divergent influences of different cerebellar manipulations (indicative of different motor diseases), the ultimate dependence of sleep architecture appears to converge on the activity of Purkinje cells. We propose that this critical node may be an ideal circuit target for therapeutic interventions in sleep disturbances linked to cerebellar dysfunction.

**Figure 6:**
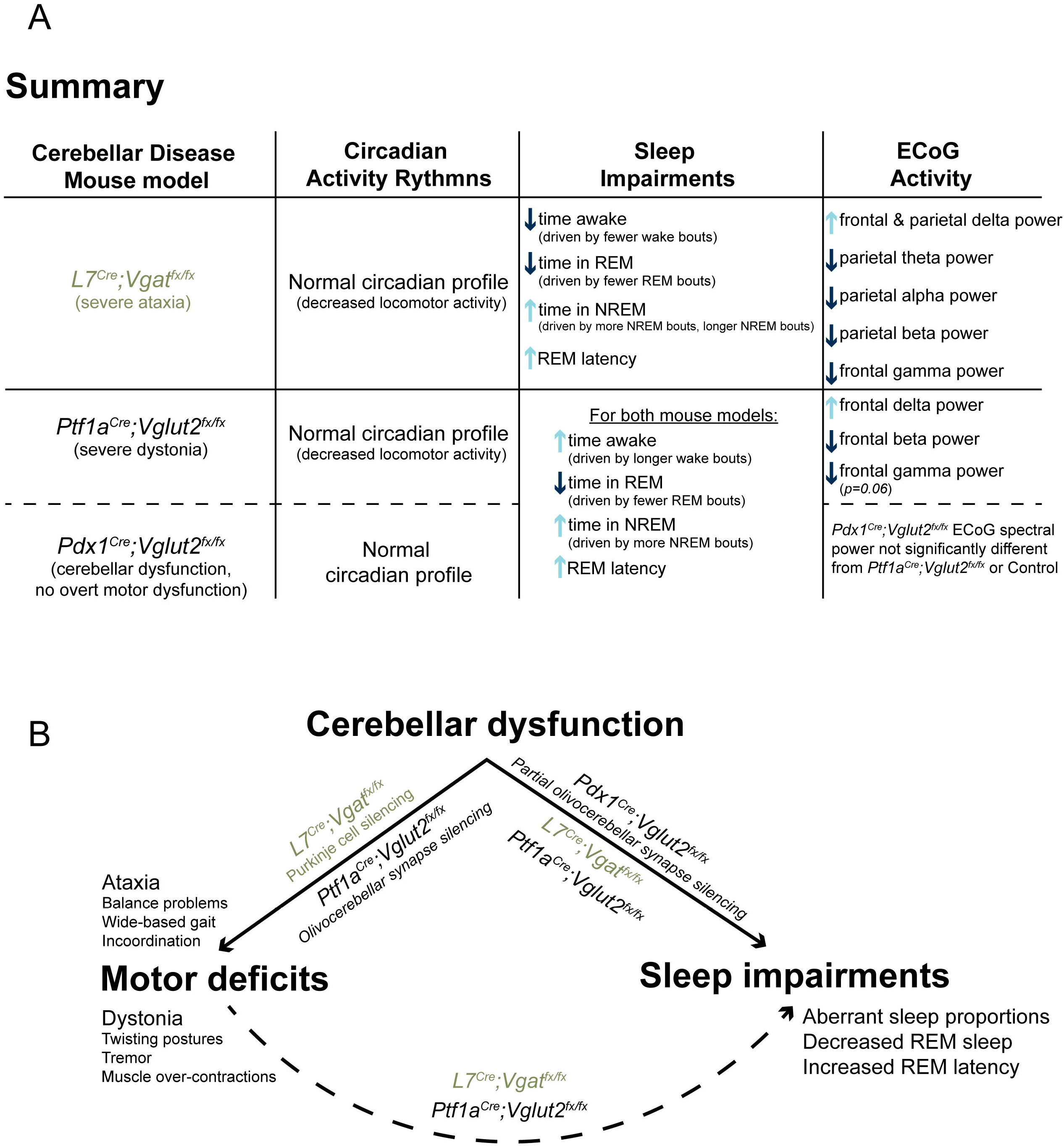
Cerebellar dysfunction causes sleep impairments across multiple motor diseases. **(A)** A summary of the main findings of this study, compared to the main findings from a complementary study in mouse models of dystonia from Salazar Leon & Sillitoe, 2023 (Leon and Sillitoe, 2023). **(B)** A proposed model for the role of the cerebellum in regulating both motor functions and specific aspects of sleep homeostasis, using evidence from multiple animal models of motor disease.

In the present study, we report decreases in total time spent in wake and REM sleep, in favor of increased time in NREM sleep in the *L7^Cre^;Vgat^fx/fx^* mouse model of ataxia. These shifted sleep proportions are driven by fewer wake and REM bouts, again in favor of a greater number of NREM bouts (Figure 4D-I, 6A). While there is limited work on sleep dysfunction in human patients with movements disorders such as ataxia, it is striking that our results recapitulate some of the patterns of sleep dysfunction observed in ataxia patients, including REM parasomnia and increased REM latency(Shindo et al., 2019; Sonni et al., 2014). Furthermore, we note that our results in *L7^Cre^;Vgat^fx/fx^* mice, particularly the deficits in REM sleep time and REM sleep latency, also recapitulate what is observed in both mouse models(Leon and Sillitoe, 2023) and human patients with dystonia(Antelmi et al., 2017; Smith, 2021), another motor disorder which involves significant cerebellar dysfunction. In this way, our work here may contribute to a growing model of cerebellar involvement in both motor and sleep dysfunction in the context of motor disorders, involving both cerebellar afferents and efferents (Figure 6B). Such a model may serve not only to contextualize the prominent sleep impairments that occur in cerebellar motor disorders, but also highlights the cerebellum as a central locus for both motor and nonmotor disease complications, which has also been discussed in other independent works(Canto et al., 2017; Salazar Leon and Sillitoe, 2022). In this context, our results may also point to the cerebellum as a key target for future therapeutics to address both the overt motor and as well the underappreciated nonmotor (sleep) dysfunctions in motor disease.

Evidence suggesting circadian dysfunction from mouse models of ataxia and human patients is mixed. While results from animal models of mild ataxia suggest that overall circadian timekeeping ability remains intact(Mendoza et al., 2010), clinical studies suggest that human patients with both hereditary spinocerebellar ataxia and idiopathic cerebellar ataxia exhibit distinct abnormalities in their circadian rhythms, including fatigue and daytime sleepiness(Sonni et al., 2014). Therefore, to clarify the role of the cerebellar circadian clock, we sought to determine whether our *L7^Cre^;Vgat^fx/fx^*mouse model of ataxia displayed any deficits in circadian timekeeping ability. Our results show that *L7^Cre^;Vgat^fx/fx^* mice display normal circadian timing of behavior, suggesting that the Purkinje cell specific manipulation is not sufficient to impact overall circadian behavior (Figure 2F-I). It is possible that, while our wheel-running measures of circadian activity did not reveal impaired circadian timekeeping ability, the decrease in awake time that we observed (Figure 3G) reflects the reported increases in fatigue and sleepiness in human ataxia patients. Additionally, while our wheel running results contrast with human patient data, it may not be surprising given the limited “access” that the cerebellum has to the circadian master clock, the suprachiasmatic nucleus (SCN). Indeed, while there are many projections between the cerebellum and other key sleep centers of the brain, including the locus coeruleus(Moises et al., 1981), pedunculopontine nucleus(Mori et al., 2016), and the hypothalamus(Dietrichs and Haines, 1989), there are no known direct projections between the cerebellum and SCN(Salazar Leon and Sillitoe, 2022; Van Dort et al., 2015). Therefore, in accordance with independent results from animal models of mild ataxia(Mendoza et al., 2010), our data suggest that the ability of the cerebellum to directly regulate circadian activity rhythms is limited. If this is the case, the fatigue and sleepiness observed in human ataxia patients may then be a result of a lack of quality sleep, peripheral mechanisms of fatigue, or even the impact of other comorbid mood disorders like depression, rather than a result of impaired circadian timekeeping ability.

Despite the emerging nature of research examining sleep dysfunction in patients with ataxia, existing studies consistently suggest a prevalence of impaired sleep, with impacts to both sleep timing and quality(Huebra et al., 2019; Patterson et al., 2018; Pedroso et al., 2011; Shindo et al., 2019; Sonni et al., 2014). We observed similar changes in arousal state dynamics and hypothesize that they are of particular interest as they may contribute to the explanation of how sleep deficits manifest in the context of motor disease. We find that *L7^Cre^;Vgat^fx/fx^* mutant mice spend less time in REM sleep and, interestingly, also spend less time awake overall compared to controls (Figure 3G-I). Instead, they spend a significantly greater amount of time in NREM sleep. One possible explanation for this disruption in sleep patterning is that Purkinje cells have been found to increase firing activity specifically during the transition from sleep to awake states(Zhang et al., 2020). As the *L7^Cre^;Vgat^fx/fx^*mouse model lacks Purkinje cell neurotransmission, it is possible that the observed decrease in wake time is a result of our genetic manipulation via the removal of the Purkinje cells’ ability to properly modulate their activity. This explanation is also parsimonious with our previous work in mouse models of dystonia which, in contrast, displayed an increase in total wake time(Leon and Sillitoe, 2023). In our dystonic mice, climbing fiber activity was genetically silenced. Previous work suggests that silencing climbing fiber activity causes an increase in Purkinje cell firing rate(Zucca et al., 2016). Therefore, it is possible that the abnormal cerebellar circuit activity in dystonia, driven by aberrant Purkinje cell activity, may be more prone to triggering the awake state while the complete silencing of Purkinje cells achieved in the *L7^Cre^;Vgat^fx/fx^* mouse model achieves this opposing dynamic of decreased total wake time (Figure 3G). Similar to humans, mouse sleep stages follow specific temporal patterns (Figure 3E) in which REM (the lightest sleep stage) is typically preceded by NREM sleep (the deepest sleep stage)(Patel et al., 2022). Importantly, this means that sleep quality is determined not only by overall time spent sleeping but also the proportions of time spent in each state. Our findings of increased NREM and decreased REM sleep time in *L7^Cre^;Vgat^fx/fx^* mice, including their overall decrease in awake time, suggest negative consequences on the quality of sleep by cerebellar dysfunction.

The identification of sleep-stage specific deficits in *L7^Cre^;Vgat^fx/fx^*mice makes sense considering the cerebellum’s established connections to and from numerous cortical regions which play roles in not only sleep regulation but also the management of specific sleep stages such as NREM and REM(Eban-Rothschild et al., 2016; Salazar Leon and Sillitoe, 2022; Van Dort et al., 2015). This observation aligns with literature suggesting that the cerebellum may regulate transitions between arousal states(Cunchillos and De Andrés, 1982; Zhang et al., 2020), and previous findings show sleep-stage specific deficits in patients with ataxia(Shindo et al., 2019; Sonni et al., 2014) and in mouse models of motor dysfunction(Leon and Sillitoe, 2023; Tsimpanouli et al., 2022). Our data showing decreased number of wake bouts (Figure 4D), decreased number of REM bouts (Figure 4E), and increased number of NREM bouts (Figure 4F) highlight and reinforce the presence of impaired sleep proportions in ataxia. These results also highlight the prevalence of REM-related sleep deficits, which we confirmed by showing an increased latency to REM sleep (Figure 4K).

Interestingly, while the changes in overall time spent in each state are reflected by the number of bouts (and for NREM, also the average bout length (Figure 4I)), the average duration of wake and REM bouts are not, as each average wake bout and REM bout is longer in the mutant mice (Figure 4G, H). It is possible that, in both cases, the elongation of bout length reflects some attempt of the brain to restore normal sleep homeostasis; as wake bouts and REM bouts are fewer in number, their length may increase to compensate. Indeed, the “REM Rebound Effect”, is common in humans and rodents following sleep deprivation or after the presence of significant stressors and involves the elongation and increased intensity of REM sleep, alongside decreased REM latency(Feriante and Singh, 2023). While the existence of a similar “Wake Rebound” is unknown, the increase in wake bout length may instead reflect the existence of some barrier to falling asleep or a decreased sleep pressure. It is known that patients with ataxia frequently report sleep-related involuntary motor behaviors like restless leg syndrome, which typically interferes with the ability to go to sleep and stay asleep(Sonni et al., 2014). It is possible a similar mechanism is responsible for the increased length of wake bouts in *L7^Cre^;Vgat^fx/fx^* mice. Alternatively, the greater number of NREM bouts may be akin to human naps, which reduce sleep pressure(Werth et al., 1996). Ultimately however, these increases in wake and REM bout length are insufficient to overcome the deficits in the number of bouts, which primarily drive the observed global impairments in sleep proportions and timing.

The increased latency to REM sleep that we observe in the *L7^Cre^;Vgat^fx/fx^*mice (Figure 4K) is particularly relevant as it reflects observations not only from patients with ataxia(Shindo et al., 2019; Sonni et al., 2014), but dystonia as well(Eichenseer et al., 2014; Smith, 2021), suggesting a role for the cerebellum in mediating REM sleep. It is possible to attribute this increased REM latency to motor dysfunction, as many REM-related sleep impairments are accompanied by involuntary motor function(Eichenseer et al., 2014; Shindo et al., 2019; Smith, 2021; Sonni et al., 2014), as the typical mechanisms of muscle atonia (a hallmark of REM sleep) are disrupted(Lydic, 2008). However, results from mouse models and humans have shown that sleep impairments in the context of motor disease can occur in the absence of motor symptoms(Antelmi et al., 2017; Leon and Sillitoe, 2023). In this case, dysfunction of the cerebellum and its circuit components may be to blame, though we acknowledge that motor dysfunction is reliably capable of causing disturbed sleep.

It is known that, of the sleep centers in the brain which project to/from the cerebellum, many are directly involved in the regulation of REM sleep(Salazar Leon and Sillitoe, 2022). In particular, regions like the locus coeruleus, which regulates NREM/REM intensity(Swift et al., 2018), sends dense projections to Purkinje cells and cerebellar nuclei neurons(Hoffer et al., 1973; Schwarz et al., 2015). Similarly the pedunculopontine nucleus, another regulator of REM sleep(Romigi et al., 2008), has afferent/efferent projections with the cerebellum, and with Purkinje cells in particular(Mori et al., 2016). Hence, the cerebellar dysfunction observed in *L7^Cre^;Vgat^fx/fx^* mice might exert a direct or indirect impact on REM latency. This could occur via intermediary regions like the pedunculopontine nucleus or the locus coeruleus, among other regions, which contribute to the regulation of REM sleep and are either a recipient or a source of direct innervation of the cerebellum(Eban-Rothschild et al., 2016; Swift et al., 2018; Van Dort et al., 2015). While the direct versus indirect circuit pathways affecting sleep regulation were not mapped in this work, our results suggest that the Purkinje-cell specific manipulation in *L7^Cre^;Vgat^fx/fx^* mice is sufficient to disrupt sleep quality, with a particular impact on REM sleep.

The observed spectral frequency oscillations in *L7^Cre^;Vgat^fx/fx^*mutant mice suggest potential network-level mechanisms underlying the altered sleep patterns (Figure 5). These changes, particularly in delta and theta bands, may raise concerns about the precision of SPINDLE, our chosen automated sleep stage scoring tool, which relies in part on frequency domain changes to score sleep stages. We address these concerns by emphasizing that SPINDLE’s algorithm extends beyond band power analysis; it incorporates multiple spectral features, assesses both ECoG and EMG activity together, and employs specific preprocessing techniques to enhance spectral pattern consistency within each recording. Furthermore, SPINDLE has been trained and validated across diverse rodent models, reinforcing its reliability amidst abnormal brain rhythms(Miladinović et al., 2019).

Our findings from the ECoG spectral activity analysis hold significant relevance in understanding the potential mechanisms of sleep deficits. We note that attributing sleep disruptions to specific changes in any frequency band poses challenges, considering both power increases and decreases across various frequency bands are linked to diverse disease states, including sleep disorders(BJORNESS et al., 2018; Long et al., 2021). Thus, we highlight the importance of recognizing spectral frequency power changes as indicative of disrupted sleep homeostasis in our ataxia mouse model, while avoiding overinterpretation of the precise directionality of these changes due to their broad implications. Still, our results may point to factors responsible for the observed disruptions in sleep and have the potential to serve as unique disease biomarkers. This insight is crucial as it helps in determining the broader impacts of cerebellar dysfunctions on brain activities, particularly in the context of sleep. For instance, we observed an increase in delta power for *L7^Cre^;Vgat^fx/fx^* mice in the parietal cortex but not frontal cortex (Figure 5C, D). Generally speaking, an increase in delta power concurs with independent work showing that higher delta power is associated with sleep impairments, particularly in instances of obstructive sleep apnea(Liu et al., 2021). This is of particular interest, as obstructive sleep apnea or general instances of sleep-disordered breathing is not only prevalent in patients with ataxia, but also thought to be regulated in part by the cerebellum(Corben et al., 2013; Kapoor and Greenough, 2015; Liu et al., 2020). Similar reductions in parietal theta power were observed (Figure 5F). The most immediate conclusion is that this reduction in theta power reflects the overall reduction in REM sleep in *L7^Cre^;Vgat^fx/fx^* mice, as it is known that theta waves are predominant during REM sleep(Merica and Blois, 1997) and often are used to characterize REM(Patel et al., 2022). Similarly, we observed a decrease in parietal alpha power (Figure 5H) in the parietal, but not frontal, cortex. We note that region-specific variations in electrographic activity are prevalent, with differential sleep-related spectral power typically observed between frontal and posterior regions of the cortex in non-disease states(Soltani et al., 2019). Therefore, our observed dichotomy of spectral power differences between parietal and frontal cortices across various frequency bands aligns with expectations. Notably, global shifts in spectral power are often associated with specific disease states, such as epilepsy(Yang et al., 2012). Thus, our observations are consistent with established patterns in the field.

It is known that feelings of sleepiness are related to a decrease in alpha power in healthy adults(Strijkstra et al., 2003). The observed decreases in parietal beta (Figure 5) and frontal gamma power (Figure 5K) also have direct associations to impairments in sleep. Beta power is typically an indicator of alert wakefulness, can be elevated in patients with primary insomnia, and decreased in patients with sleep impairments like obstructive sleep apnea(Liu et al., 2021). Though assessment of sleep apnea is beyond the scope of this work, the observed decrease in beta power in *L7^Cre^;Vgat^fx/fx^*mice agrees with the role of beta waves as an indication of being awake, as *L7^Cre^;Vgat^fx/fx^* mice appear to spend less time awake. Interestingly, bursts of beta activity can also occur during REM sleep(Steriade, 2009). As total REM time is lower in the mutant mice, this may also explain the observed decrease in beta power. Similarly, gamma power can indicate working memory and attention, which is expected to decrease in *L7^Cre^;Vgat^fx/fx^*mice that display decreased awake time(Goddard et al., 2012). Spontaneous gamma activity also occurs during REM sleep(Steriade, 2009); the decreased gamma power may then reflect the same decrease in REM. Overall in this work, we demonstrate that precise genetic manipulation of a single cerebellar cell type leading to motor deficits also disrupts arousal state dynamics, implying a necessary role for cerebellar Purkinje cells in maintaining normal sleep patterns and contributing to their disruption in the context of ataxia.

## Conclusion

By exploiting the genetic precision of the *L7^Cre^;Vgat^fx/fx^*mouse circuit model of ataxia, we tested if sleep dysregulation depends on cerebellar Purkinje cells. Our data supports the possibility of a critical role for the cerebellum in sleep regulation, which is reflected in the patterns of sleep disruption that are observed in human movement disorders. These findings not only expand our understanding of the cerebellum’s involvement in nonmotor complications within motor diseases but also suggest that cerebellar circuitry could drive similar sleep deficits across different motor disorders (Figure 6B). This knowledge points to potential broader network dysfunction in motor disorders, with the cerebellum poised at the nexus of various disease symptoms. Consequently, the insights from our model illuminate the relevance of sleep impairments in human cerebellar motor disorders and position the cerebellum as a promising target for therapy.

## Materials and Methods

### Ethics

Animal experimentation: All animals were housed in an AALAS-certified facility that operates on a 14-hour light cycle. Husbandry, housing, euthanasia, and experimental guidelines were reviewed and approved by the Institutional Animal Care and Use Committee (IACUC) of Baylor College of Medicine (protocol number: AN-5996).

### Animals

All mice used in this study were housed in a Level 3, AALAS-certified facility. All experiments and studies that involved mice were reviewed and approved by the Institutional Animal Care and Use Committee of Baylor College of Medicine (BCM AN-5996). We purchased *L7^Cre^*(L7Cre-2, #004146)(Lewis et al., 2004) and *Vgat*-floxed (*Vgat^flox^*, #012897)(Tong et al., 2008) mice from The Jackson Laboratory (Bar Harbor, ME, USA) and then maintained them in our colony using a standard breeding scheme. The conditional knock-out mice that resulted in ataxia were generated by crossing *L7^Cre^;Vgat^fx/fx^* heterozygote mice with homozygote *Vgat^fx/fx^* mice. *L7^Cre^;Vgat^fx/fx^* mice were considered experimental animals. A full description of the genotyping details (e.g., primer sequences and the use of a standard polymerase chain reaction) was provided in White et al., 2014 (White et al., 2014). All littermates lacking *Cre* upon genotyping were considered control mice. Ear punches were collected before weaning and used for genotyping and identification of the different alleles. For all experiments, we bred mice using standard timed pregnancies, noon on the day a vaginal plug was detected was considered embryonic day (E)0.5 and postnatal day (P)0 was defined as the day of birth. Mice of both sexes were used in all experiments.

### Tissue preparation and processing for *in situ* hybridization

Mice were anaesthetized with isoflurane and brains were removed from the skull and immersed in OCT (optimal cutting temperature). Immersed brains were flash frozen by placing tissue molds onto dry ice. Sagittal sections (25 μm) were cut through the cerebellum and the slices placed onto electrostatically coated glass slides (Probe On Plus Fisher Brand; Fisher Scientific). The tissue was probed with *Cre* or *Vgat* (SLC32A1) digoxigenin-labelled mRNA probes using an automated in situ hybridization procedure (Genepaint). All reagent incubations, washes and stains were automated and performed by the *in situ* hybridization robot. The signal was detected by colorimetric detection using BCPI/NBT reagents. After processing was completed, the slides were removed from the machine and then cover-slipped with permanent mounting medium (Entellan mounting media, Electron Microscopy Sciences, Hatfield, PA, USA) and left to dry before imaging.

### Wheel-running behavior

Recordings were maintained in a ventilated, temperature-controlled, and light-tight room under either a 12:12 LD cycle or DD conditions. Mice were singly housed in wheel-running cages and allowed to entrain to the LD cycle for two weeks, before being released into DD conditions for 21 days, to assess endogenous circadian timekeeping ability. We assessed period length, activity onset, phase shift onset, and average number of wheel revolutions per five minutes using ClockLab Analysis (Actimetrics). All measures were calculated automatically by the Clocklab Analysis software.

### Surgical procedure for ECoG/EMG sleep recordings

Prior to surgery, mice were given preemptive analgesics (extended-release buprenorphine, 1 mg/kg subcutaneous (SC), and meloxicam, 5 mg/kg SC) with continued application as part of the standard 3 day post-operative procedure. Mice were anesthetized with isoflurane and placed into a stereotaxic device, which continued to deliver isoflurane throughout surgery. Each mouse with implanted with a prefabricated ECoG/EMG headmount (Pinnacle Technology, Lawrence KS, #8201) with 0.10” EEG screws to secure headmounts to the skull (Pinnacle Technology, Lawrence KS, #8209). To do this, fur was removed with depilatory cream (Nair) and the surgical site was sterilized with alternating applications of alcohol and betadine scrub solution. Then a midline incision was made, and the skull was exposed. The headmount was affixed to the skull using cyanoacrylate glue to hold it in place while pilot holes for screws were made and screws were inserted. Screws were placed bilaterally over parietal cortex and frontal cortex. A small amount of silver epoxy (Pinnacle Technology, Lawrence KS, #8226) was applied to the screw-headmount connection. Platinum-iridium EMG wires on the prefabricated headmount were placed under the skin of the neck, resting directly on the trapezius muscles. The headmount was permanently affixed to the skull using ‘Cold-Cure’ dental cement (A-M systems, #525000 and #526000). Mice were allowed to recover for three to four days before being fitted with a preamplifier (Pinnacle Technology, Lawrence KS, #8202) and tethered to the recording device (Pinnacle Technology, Lawrence KS, #8204 and #8206-HR).

### ECoG/EMG sleep recordings

Mice were recorded in light and temperature-controlled rooms, for eight hours, at the same time of day for every mouse. The first hour of recording was considered the acclimation period and was therefore excluded from final analysis. Food and water were available *ad libitum* throughout the recording day. Mice were singly housed in clear acrylic cages (Pinnacle Technology, Lawrence KS, #8228). Preamplifiers were connected to a 360° commutator allowing for unrestricted movement (Pinnacle Technology, Lawrence KS, #8204). All data was collected by the Data Acquisition and Conditioning System (Pinnacle Technology, Lawrence KS, #8206-HR) which was specifically tuned for detecting sleep. Data were captured using Sirenia Acquisition software (Pinnacle Technology, Lawrence KS).

### Sleep scoring and analysis of sleep data

Sleep was automatically scored offline via SPINDLE(Miladinović et al., 2019). For spectral frequency analysis of ECoG and EMG activity, raw files were also pre-processed in MATLAB (MathWorks) using the free toolkit EEGLAB (UC San Diego). Scored files were downloaded from SPINDLE as a .csv and statistical analysis was performed in R v4.1.2.

### Data analysis and statistics

Data are presented as mean ± SEM and analyzed as a Two Sample t-test (normally distributed data with equal variance), Welch Two Sample t-test (normally distributed data with unequal variance), or Wilcoxon rank sum exact test (non-normally distributed data with unequal variance). For all statistical tests, p < 0.05 was considered as statistically significant. All statistical analyses were performed using R v4.1.2.

## Acknowledgements and Funding Sources

This work was supported by Baylor College of Medicine (BCM), Texas Children’s Hospital, The Hamill Foundation, and the National Institutes of Neurological Disorders and Stroke (NINDS) R01NS100874, R01NS119301, and R01NS127435 to RVS. Research reported in this publication was supported by the Eunice Kennedy Shriver National Institute of Child Health & Human Development of the National Institutes of Health under Award Number P50HD103555 for use of the Animal Behavior Core and the Cell and Tissue Pathogenesis Core (BCM IDDRC). The content is solely the responsibility of the authors and does not necessarily represent the official views of the National Institutes of Health. Support was also provided by a Dystonia Medical Research Foundation (DMRF) grant to RVS.

Research reported in this publication was supported in part by the RNA *In Situ* Hybridization Core facility at Baylor College of Medicine, which is supported by a Shared Instrumentation grant from the NIH (S10OD016167) and the NIH IDDRC grant P50HD103555 from the Eunice Kennedy Shriver National Institute of Child Health & Human Development. The content is solely the responsibility of the authors and does not necessarily represent the official views of the Eunice Kennedy Shriver National Institute of Child Health & Human Development or the National Institutes of Health.

## Author contributions

Technical and conceptual ideas in this work were conceived by LESL and RVS. LESL, AMB, and HK performed experiments. LESL and RVS performed data analysis and data interpretation. LESL and RVS wrote the manuscript. LESL, AMB, HK, and RVS edited the manuscript.

## Data availability

All data generated or analyzed in this study are included in the manuscript and supporting files.

